# Providing an additional electron sink by the introduction of cyanobacterial flavodiirons enhances the growth of *Arabidopsis thaliana* in varying light

**DOI:** 10.1101/2020.02.05.935346

**Authors:** Suresh Tula, Fahimeh Shahinnia, Michael Melzer, Twan Rutten, Rodrigo Gómez, Anabella F. Lodeyro, Nicolaus von Wirén, Néstor Carrillo, Mohammad R. Hajirezaei

## Abstract

The ability of plants to maintain photosynthesis in a dynamically changing environment is of central importance for their growth. As their photosynthetic machinery typically cannot adapt rapidly to fluctuations in the intensity of radiation, the level of photosynthetic efficiency is not always optimal. Cyanobacteria, algae, non-vascular plants (mosses and liverworts) and gymnosperms all produce flavodiirons (Flvs), a class of proteins not represented in the angiosperms; these proteins act to mitigate the photoinhibition of photosystem I. Here, genes specifying two cyanobacterial Flvs have been expressed in the chloroplasts of *Arabidopsis thaliana* in an attempt to improve the robustness of Photosystem I (PSI). The expression of *Flv1* and *Flv3* together shown to enhance the efficiency of the utilization of light and to boost the plant’s capacity to accumulate biomass. Based on an assessment of the chlorophyll fluorescence in the transgenic plants, the implication was that photosynthetic activity (including electron transport flow and non-photochemical quenching during a dark-to-light transition) was initiated earlier in the transgenic than in wild type plants. The improved photosynthetic performance of the transgenics was accompanied by an increased production of ATP, an acceleration of carbohydrate metabolism and a more pronounced partitioning of sucrose into starch. The indications are that Flvs are able to establish an efficient electron sink downstream of PSI, thereby ensuring that the photosynthetic electron transport chain remains in a more oxidized state. The expression of Flvs in a plant acts to both protect photosynthesis and to control the ATP/NADPH ratio; together, their presence is beneficial for the plant’s growth potential.

## Introduction

The growth and development of plants, fueled by photosynthesis, depends on the capture of light energy (Stitt et al., 2010), a process which is carried out by the chloroplast. Photosynthesis can be inhibited by either an inadequate pool of ATP or an imbalance between the quantity of ATP and NADPH present (Avenson et al., 2005b; Cruz et al., 2005; Amthor, 2010). Adjustments of ATP in chloroplasts through CET pathway results in the non-photochemical quenching (NPQ) of Photosystem II (PSII), resulting in a significant loss in the fixation of CO_2_ (Long et al., 1994, Werner et al., 2001, Zhu et al., 2004). Fine tuning of NPQ in tobacco has enabled an enhanced level of photosynthetic efficiency to be attained, leading to a measurable increase in biomass yield (Kromdijk et al., 2016). One of the consequences of reducing photosynthetic efficiency is the accumulation of surplus electrons in the photosynthetic electron transport chain (PETC), which promotes the production of reactive oxygen species (ROS). Since the functioning of both PSI and PSII is compromised by the presence of ROS (Miyake, 2010), the result is a diminution in the plant’s capacity to assimilate CO_2_ (Zivcak et al., 2015a,b; Takagi et al., 2016). According to Gollan et al. (2017), the inhibition of PSI experienced by the *Arabidopsis thaliana pgr5* mutant when exposed to a high light intensity limits both the light and dark reactions associated with photosynthesis, as well as chloroplast signaling and nuclear gene expression. Thus, avoiding PSI electron acceptor limitation or introducing additional electron dissipating pathways into the chloroplast have the potential to improve photosynthetic efficiency and hence increase the plant’s productivity.

Algae, cyanobacteria, non-vascular plants (mosses and liverworts) and gymnosperms have evolved an alternative electron flow (AEF) pathway, driven by the so-called flavodiirons (Flvs). Analysis of the genome of the cyanobacterium *Synechocystis* sp. PCC6803 has identified the presence of four *Flv* genes. Their products, in the form of the heterodimers Flv1/Flv3 and Flv2/Flv4 drive oxygen-dependent electron flow under low levels of CO_2_ availability and fluctuating light conditions (Zhang et al., 2012; Allahverdiyeva et al., 2013; Hayashi et al., 2014; Shimakawa et al., 2015). The Flv1/Flv3 heterodimer generates an electron sink downstream of PSI and directs the electron flow to reduce O_2_ to H_2_O, thereby protecting PSI (Allahverdiyeva et al., 2013). Cyanobacterial Flv3 can also form a functional homodimer, which in *Chlamydomonas reinhardtii* acts in conjunction with ferredoxin 9 to transfer electrons into a glutathione-cysteine peroxidase pathway (Cassier-Chauvat et al., 2014). According to Yamamoto et al. (2016), inducing the expression of the *Physcomitrella patens* Flv1/Flv3 heterodimer in the *A. thaliana pgr5* mutant provides partial compensation for the impairment of the cyclic electron transport pathway, an observation which demonstrates that Flvs can be functionally beneficial in an angiosperm. Shimakawa et al. (2017) have shown that in the liverwort *Marchantia polymorpha*, Flv1 contributes to P700 oxidation and hence protects PSI against photoinhibition. When Gómez et al. (2018) expressed the cyanobacterial genes *Flv1* and *Flv3* in tobacco chloroplasts, the photosynthetic performance under steady-state illumination of the resulting transgenic plants proved to be comparable to that of wild type (WT) leaves, while the recovery of electron transport and NPQ during a dark-to-light transition was more rapid. The joint expression *Flv1* and *Flv3* isolated from a moss species in either the rice mutants *pgr5* or *NDH* has been shown to rescue biomass accumulation (Wada et al., 2017). Meanwhile, the enhancement to ATP synthesis resulting from the over-expression of *Flv3* in *Synechocystis* results in an accumulation of glycogen and a consequent increase in cell dry weight (Hasunuma et al., 2014).

The purpose of the present investigation was to determine the phenotypic effect of expressing *Synechocystis Flv* genes in *A. thaliana*. The focus was to establish whether heterologously expressed Flv could mediate electron transfer to oxygen within the PETC, and if so, whether this capacity had the potential to boost the plant’s productivity.

## Materials and methods

### The heterologous expression of cyanobacterial Flvs and the localization of transgene products

*A. thaliana* Col-0 lines constitutively expressing either *Flv1/Flv3* were generated using the floral dip method (Clough et al., 1998), using the pCHF3-derived vectors described by Gómez et al. (2018). Transgene expression was driven by the cauliflower mosaic virus (CaMV) 35S promoter, and a pea sequence encoding a ferredoxin-NADP^+^ reductase transit peptide was fused in-frame to the 5’ end of the transgene to direct its expression to the chloroplast (Figure S1a). Five independent transformants were chosen for each construct, and the presence of the transgene in the host’s nuclear genome was confirmed by PCR analysis. Of the five transformants selected per construct, three were developed into fixed transgenic lines: their ability to accumulate *Flv* transcript was monitored using a quantitative real time PCR (qRT-PCR) assay (see below). The level of transcription of the various transgenes is shown in Figure S1b. The chloroplast targeting of the transgenes was validated by fusing the *GFP* sequence (encoding green fluorescent protein) to the C terminus of the *Flv* transgene, taking advantage of PGBW5 Gateway binary vectors driven by the 35S CaMV promoter (Figure S2). The binary vector containing the *Flv* transgene was transferred into *Agrobacterium tumefaciens* strain EHA 105 using the Dower et al. (1988) electroporation method, and from thence into leaves of *Nicotiana benthamiana* using the agroinfiltration method described by Sainsbury and Lomonossoff (2008). Leaves sampled 48 h after infiltration was subjected to confocal scanning microscopy to monitor GFP activity (Figure S2).

### Experimental growing conditions for A. thaliana transgenics

Seed of each of the *Flv* transgenic lines and WT were surface-sterilized by immersion for 15 min in 70% (v/v) ethanol and 0.05% (v/v) Tween 20, then rinsed for 30 s in 96% (v/v) ethanol. After air drying, the seed was held at 4°C for 48 h, plated on vertically-orientated agar containing half strength Murashige and Skoog (1962) medium and held under an 8 h photoperiod (160 µmol photons m^-2^ s^-1^) at 22°C. After two weeks, the seedlings were potted into a mixture of 70 L substrate 1 (Duesseldorf, Germany), 23 L vermiculite, 372 g plantacote depot 4m and held at 22°C, 80% relative humidity under an 8 h photoperiod provided by Master HPI-T Plus 250 W fluorescent lights (Philips, Netherlands) involving four different light intensities: low (50 µmol photons m^-2^ s^-1^), moderate (160 µmol photons m^-2^ s^-1^), moderately high (300 µmol photons m^-2^ s^- 1^) and high (600 µmol photons m^-2^ s^-1^). The CO_2_ level was maintained at 400 ppm and plants were kept fully hydrated. Shoot dry weights were derived from six week old plants. For the biochemical analysis of leaf, rosettes of six week old plants exposed to 160 µmol photons m^-2^ s^-1^ harvested at various time points during the diurnal cycle (0, 4, 8, 16, 20 and 24 h) were snap-frozen in liquid nitrogen and ground to a powder. A number of plants were also grown under long day conditions (16 h photoperiod, 160 µmol photons m^-2^ s^-1^), with all the other environmental parameters identical with those used for the short day grown plants.

### Determination of the leaf content of carbohydrates, amino acids and metabolites

For the determination of the content of soluble sugars (glucose, fructose and sucrose) and amino acids, a 50 mg aliquot of powdered frozen leaf was extracted in 0.7 mL 80% (v/v) ethanol at 80°C for 1 h. Following centrifugation (14,000 rpm, 10 min), the supernatant was evaporated under vacuum at 40°C, and the residue dissolved in 0.2 mL deionized water. Sugar contents were quantified using the enzymatic method described by Ahkami et al. (2013), while those of the individual amino acids were obtained following Mayta et al. (2018). The pelleted material was used to assess the leaf’s starch content: the pellet was rinsed twice in 80% (v/v) ethanol, air-dried at 80°C for 1 h and resuspended in 0.2 M KOH. The resulting suspension was held at 80°C for 1 h, the pH adjusted to neutrality using 1 M acetic acid, then incubated overnight at 37°C in 50 mM NaAc (pH 5.2) containing 7 units mg^-1^ amyloglucosidase. The glucose thereby released was quantified as above. The methods used for the quantification of primary metabolites followed Ghaffari et al. (2016).

### Determination of the leaf content of adenine phosphate and redox equivalents

Adenine phosphates were quantified employing a UPLC-based method developed from that described by Haink and Deussen (2003). Prior to the HPLC separation step, a 20 µL aliquot of the sample used for the quantification of metabolites (as well as a mixture of ATP, ADP, AMP and ADPGlc) were derivatized by the addition of 45 µL 10% (v/v) chloracetaldehyde and 435 µL 62 mM sodium citrate/76 mM KH_2_PO_4_ (pH 5.2), followed by a 40 min hold at 80°C, cooling on ice, and a centrifugation (20,000 *g*, 1 min). Reverse phase UPLC separations were achieved using an Infinity 1200 device (Agilent, Waldbronn, Germany). The gradient was established using eluents A [TBAS/KH_2_PO_4_ (5.7 mM tetrabutylammonium bisulfate/30.5 mM KH_2_PO_4_, pH 5.8)] and B (a 2:1 mixture of acetonitrile and TBAS/KH_2_PO_4_); the Roti C Solv HPLC reagents were purchased from Roth (Karlsruhe, Germany). The 1.8 µm, 2.1×50 mm separation column was an Eclipse plus C18. The column was pre-equilibrated for at least 30 min in a 9:1 mixture of eluents A and B; during the first two minutes of the run, the column contained 9:1 A:B, changed thereafter to 2:3 A:B for 2.3 minutes followed by a change to 1:9 A:B for 3.1 minutes and set to initial values of 1:9 for 2.6 minutes. The flow rate was 0.6 mL min^-1^ and the column temperature was maintained at 37°C. The excitation and emission wavelengths were, respectively 280 nm and 410 nm. Chromatograms were integrated using MassHunter (release B.04.00) software (Agilent, Waldbronn, Germany).

For the quantification of the redox equivalents, a 0.5 g sample of powdered leaf was suspended in 1 ml 0.1 M KOH dissolved in 50% (v/v) ethanol (NADPH, NADH) or in 1 mL 0.1 M HCl (NADP^+^, NAD^+^). The entire extraction procedure was carried out under green light to minimize degradation of the analytes. The extractions were held on ice for 5 min, then centrifuged (14,000 rpm, 10 min, 4°C) and the supernatants subjected to ion chromatography coupled MS/MS following Ghaffari et al. (2016). However, the recovery experiment of NADPH and NADH in the samples was 50%. The results after light for 4 h could not be reliably calculated since the values were bellow detection limit and the results after light for 8 h did not show significant differences between the WT and different transgenic plants (Figure S3).

### Determination of the leaf content of glutathione

The method used to extract glutathione from the leaf material followed Davey et al. (2003). The separation and detection methods were established as follow. Approximately 100 mg of fresh leaf material was ground to fine powder using tissue homogenizer with 1 mM EDTA and 0.1% (v/v) formic acid at 4°C under green safe light and centrifuged at maximum speed (35,280 *g*) for 10 min. Measurements of oxidized and reduced glutathione were carried out immediately in freshly prepared extracts. Separation and analysis of the desired compounds was performed out on a C18 column (HSS T3, 1.8 µm, 2.1×150 mm, Waters Germany) and an UPLC/MS-MS (Infinity ll, 6490 Triple Quadrupole LC/MS, Agilent, Waldbronn, Germany), respectively. Two µL of extracts and the corresponding standards were injected in the mobile phase consisting of purest water plus 0.1% (v/v) formic acid and pure methanol plus 0.1% (v/v) formic acid. The temperatures of the auto sampler and column were maintained at 8°C and 37°C, respectively. Separated compounds were eluted at a flow rate of 0.5 mL min^-1^. The evaluation and quantification of the compounds were performed using the software MassHunter, release B.04.00 (Agilent, Waldbronn, Germany).

### Determination of photosynthetic parameters

The quantification of chlorophyll *a* was based on fluorescence measurements obtained using a multispeQ v1.0 device (photosynq.org) in conjunction with photosynQ software (Kuhlgert et al., 2016). Measurements were taken from the fully expanded leaves of six week old plants held in the dark for 4 h. The minimal fluorescence parameter *F*_0_ was measured before exposing the leaf to a pulse of red (650 nm) light (500 ms, 3,000 μmol photon m^−2^ s^−1^), from which the maximal fluorescence parameter (*F*_m_) was obtained. *F*_*m*_′ and *F*_*s*_ were determined during exposure to 160 μmol photon m^−2^ s^−1^ actinic light. Chlorophyll transient induction measurements on dark-adapted leaves were carried out following Gómez et al. (2018). PSII parameters were calculated based on the suggestion of Baker (2008), namely: *F*_v_/*F*_m_=(*F*_m_–*F*_o_)/*F*_m_ and NPQ=(*F*_m_–*F*_m_′)/*F*_m_′. The parameters Φ_PSII,_ Φ_NPQ,_ Φ_No_ and qL were calculated according to Kramer et al. (2004).

### Transmission electron and confocal laser scanning microscopy

Transmission electron microscopy was performed following Mayta et al. (2018). A 2 mm^2^ piece cut from the center of the tobacco leaf was taken from both WT and *Flv* transgenics, and processed according to Kraner et al. (2017). For confocal laser scanning microscopy, the leaf samples were degassed to remove any entrapped air, and GFP activity was detected using an LSM780 device (Carl Zeiss, Jena Germany), with the excitation wavelength set to 488 nm; fluorescence was detected over the wavelength range 491-535 nm.

### RNA isolation, cDNA synthesis and transcription analysis

Total RNA was extracted from young leaves following the protocol given by Logemann et al. (1987) after which it was subjected to DNase treatment (Life Technologies, Darmstadt, Germany) and converted to ss cDNA using a RevertAid first strand cDNA synthesis kit (Life Technologies, Darmstadt, Germany) provided with a template of 1 μg total RNA and oligo dT primer: the reaction was run at 42°C for 60 min. The primers used for qRT-PCR analysis of *Flv* transgenes are given in Table S4. The assays were performed in a CFX384 touch real-time system using the SYBR Green Master Mix Kit (Bio-Rad, Feldkirchen, Germany). The primer pairs employed to amplify *Flv1* (Flv1-RT F/R) and *Flv3* (Flv3-RT F/R), along with those amplifying the reference sequence *Ubi10* (GenBank accession number *At4g05320*) are listed in Table S4. Relative transcript abundances were determined using the method Schmittgen and Livak (2008) method.

### Statistical analysis

Means and standard errors (SE) were calculated using SigmaPlot software (www.sigmaplot.co.uk/products/sigmaplot/sigmaplot-details.php). The Students’ *t*-test was used to test for the statistical significance of differences between means.

## Results

### The growth response of Flv transgenics to variation in the light intensity

The development of biomass in both WT and *Flv* transgenic plants grown at varying light intensities is illustrated in Figure 1. When illuminated with 50 µmol photons m^-2^ s^-1^, the performance of the transgenic plants was not distinguishable from that of the WT ones. However, when the intensity was increased to either 160 or 300 µmol photons m^-2^ s^-1^, the transgenic plants were clearly larger (Figure 1a), as confirmed by comparisons of their shoot dry weight in transgenic plants harbored *Flv1*/*Flv3*, out-performed WT by 10-30% (Figure 1b). Among the plants exposed to the highest intensity (600 µmol photons m^-2^ s^-1^), the only lines which were superior to WT were those expressing *Flv1*/*Flv3*.

**Figure 1.**
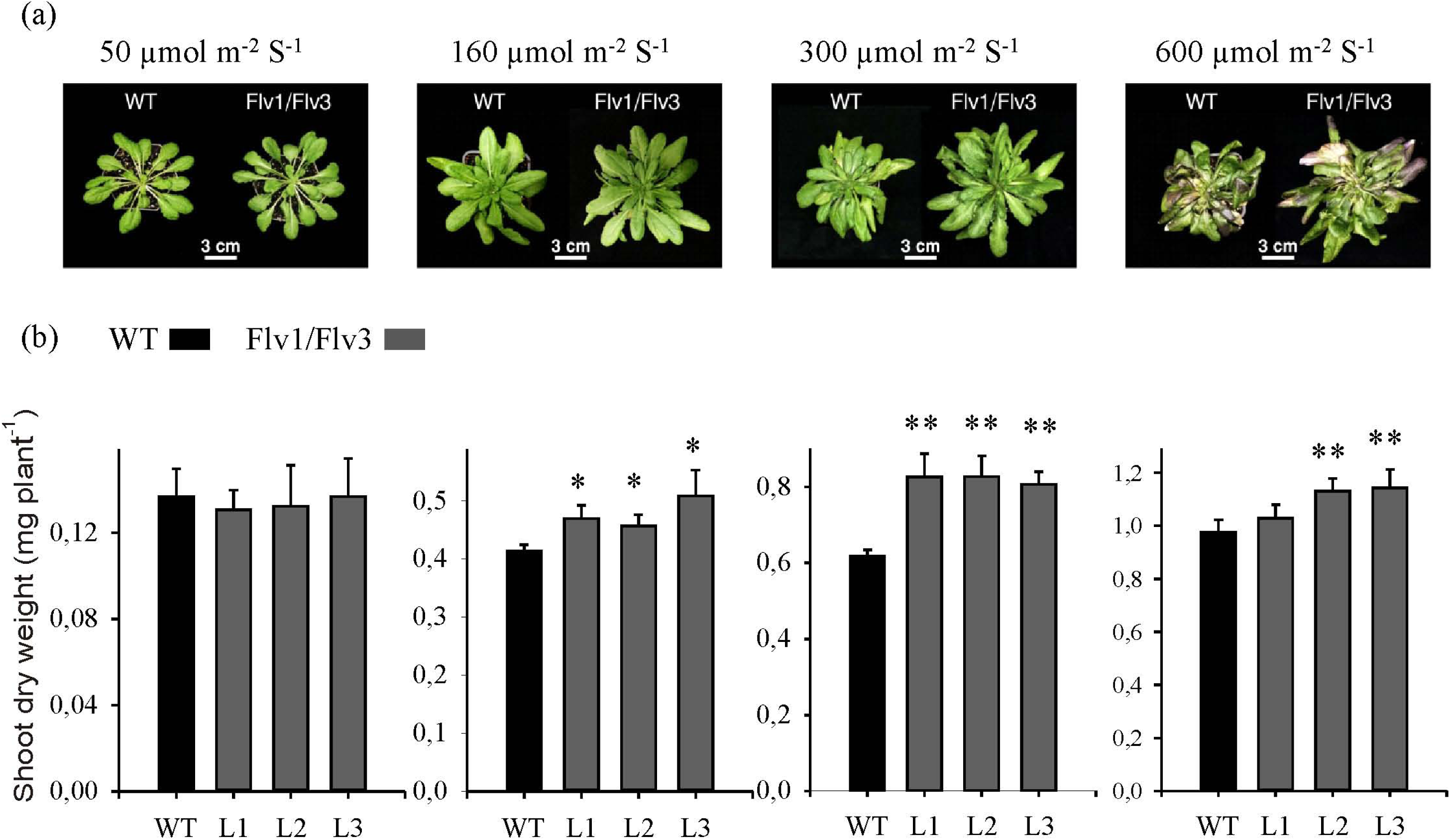
The growth of *A. thaliana* plants heterologously expressing cyanobacterial *Flv* genes. (a) The phenotype of six week old plants exposed to an 8 h photoperiod at a variety of light intensities. (b) Shoot dry weight: data are given in the form mean±SE (*n=*5-8). *,**: means differ from the performance of WT at, respectively, P≤0.05 and ≤0.01.

### The effect of expressing Flv transgenes on the biomass accumulation of plants exposed to long days

Under a long day regime, the *Flv* transgenics reached flowering earlier than WT plants (data not shown), and were more bushy, with increased inflorescences (Figure 2a). Their shoots dry weight was up to 1.8 fold greater (Figure 2b). Seed size was unaffected by the presence of the transgene (data not shown), but seed yield was 1.2-1.8 fold greater (Figure 2c).

**Figure 2.**
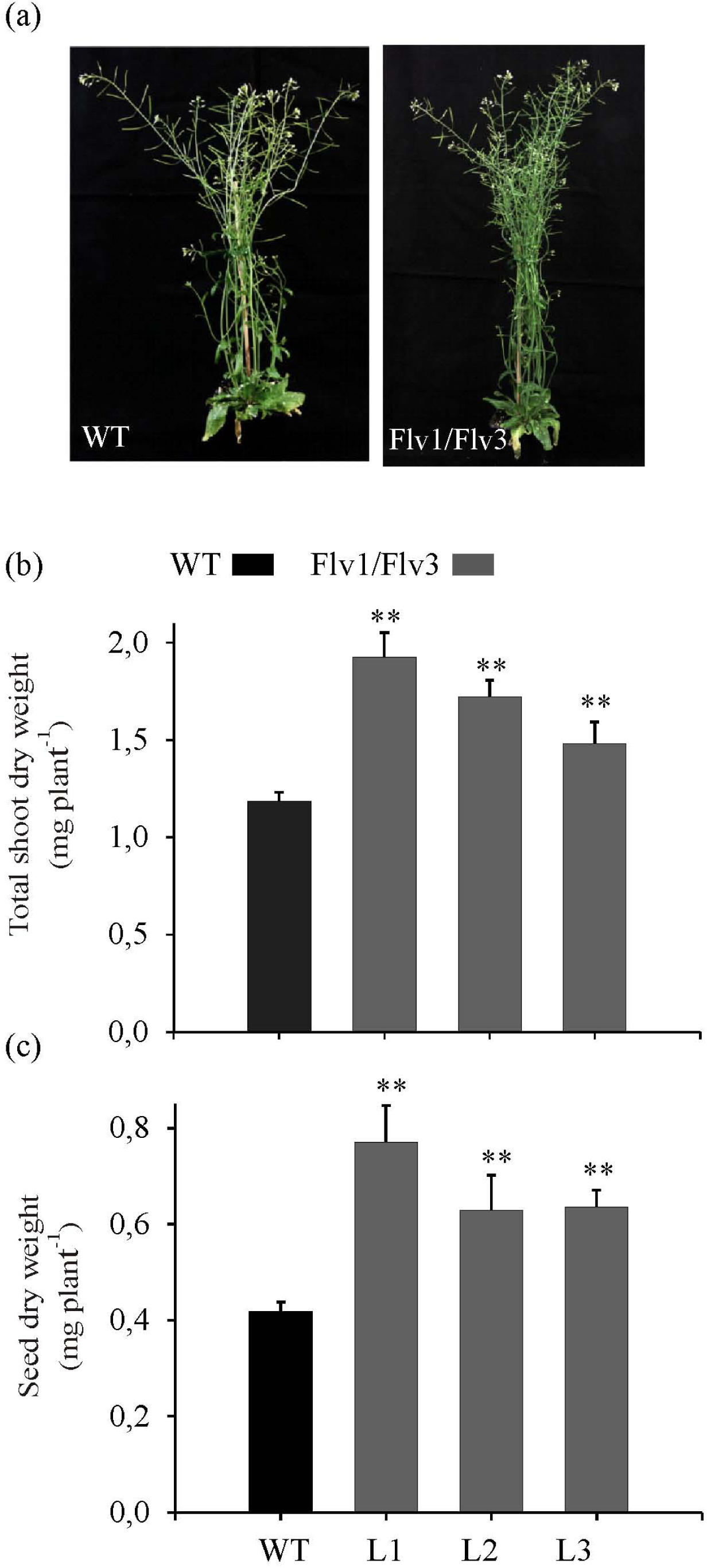
The effect of heterologously expressing *Flv* genes on the phenotype of long day grown plants. (a) Fully grown WT and *Flv* transgenics grown under a 16 h photoperiod (160160 μmol photons m^−2^ s^−1^ of actinic light). (b) Shoot dry weight, (c) seed yield per plant. Data shown in the form mean±SE (*n=*8). **: means differ from the performance of WT at ≤0.01.

### Chlorophyll fluorescence in the leaf of Flv transgenics

Chlorophyll fluorescence measurements were taken from the fully expanded leaves of dark-adapted six week old plants. The F_v_/F_m_ ratio (a measure of the maximum quantum yield of PSII) was somewhat higher in the leaves of the *Flv1/Flv3* transgenic than in those of WT (Figure 3a). In leaves exposed to 160 µmol photons m^-2^ s^-1^, Φ_PSII_ (representing the quantum yield of PSII) in the *Flv1/Flv3* transgenics showed higher than in WT leaves (Figure 3b). Similarly, qL (an estimate of the fraction of open reaction centers) was greater in the leaves of the *Flv1/Flv3* transgenics than in WT leaves (Figure 3c). Φ_NPQ_ (the quantum yield of quenching due to light-induced processes) was significantly lower during the measurement in the *Flv1/Flv3* trangenics than in WT (Figure 4a). In contrast, Φ_NO_ (the quantum yield of non-light-induced quenching processes) in *Flv1/Flv3* trangenics clearly shows no difference to the WT level (Figure 4b). Finally, NPQ (non-photochemical quenching) was up to two fold times higher than the WT level in the *Flv1*/*Flv3* transgenics (Figure 4c).

**Figure 3.**
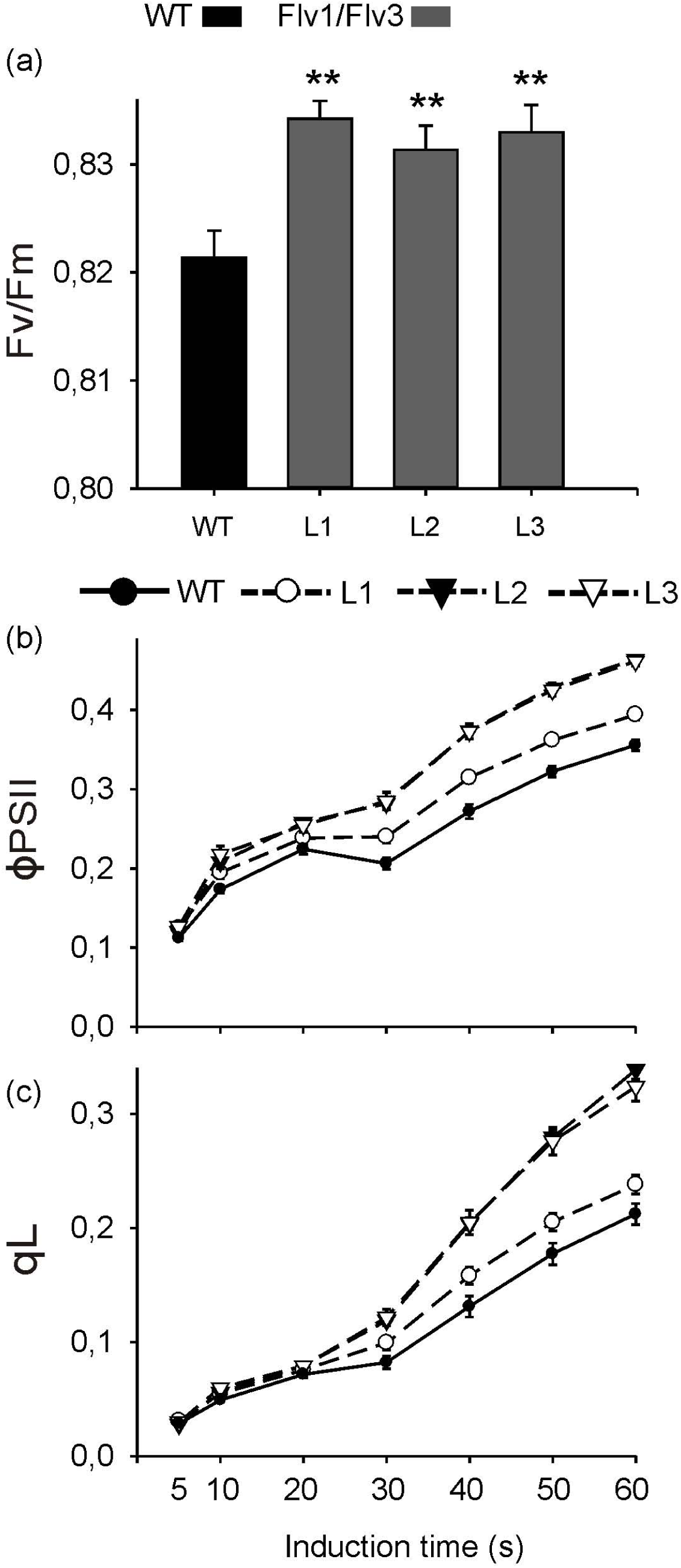
The effect of heterologously expressing *Flv* genes on PSII activity and the oxidation state of the electron transport chain. L1-L3: lines harboring *Flv1/Flv3.* (a) F_v_/F_m_ (the maximum quantum yield of PSII). **: means differ from the performance of WT at ≤0.01. The temporal behavior of dark-adapted leaves of six week old plants exposed to 160 μmol photons m^−2^ s^−1^ of actinic light with respect to (b) Φ_PSII_ (the quantum yield of PSII), (c) qL (the fraction of open PSII reaction centers). Data shown in the form mean±SE (*n=*8).

**Figure 4.**
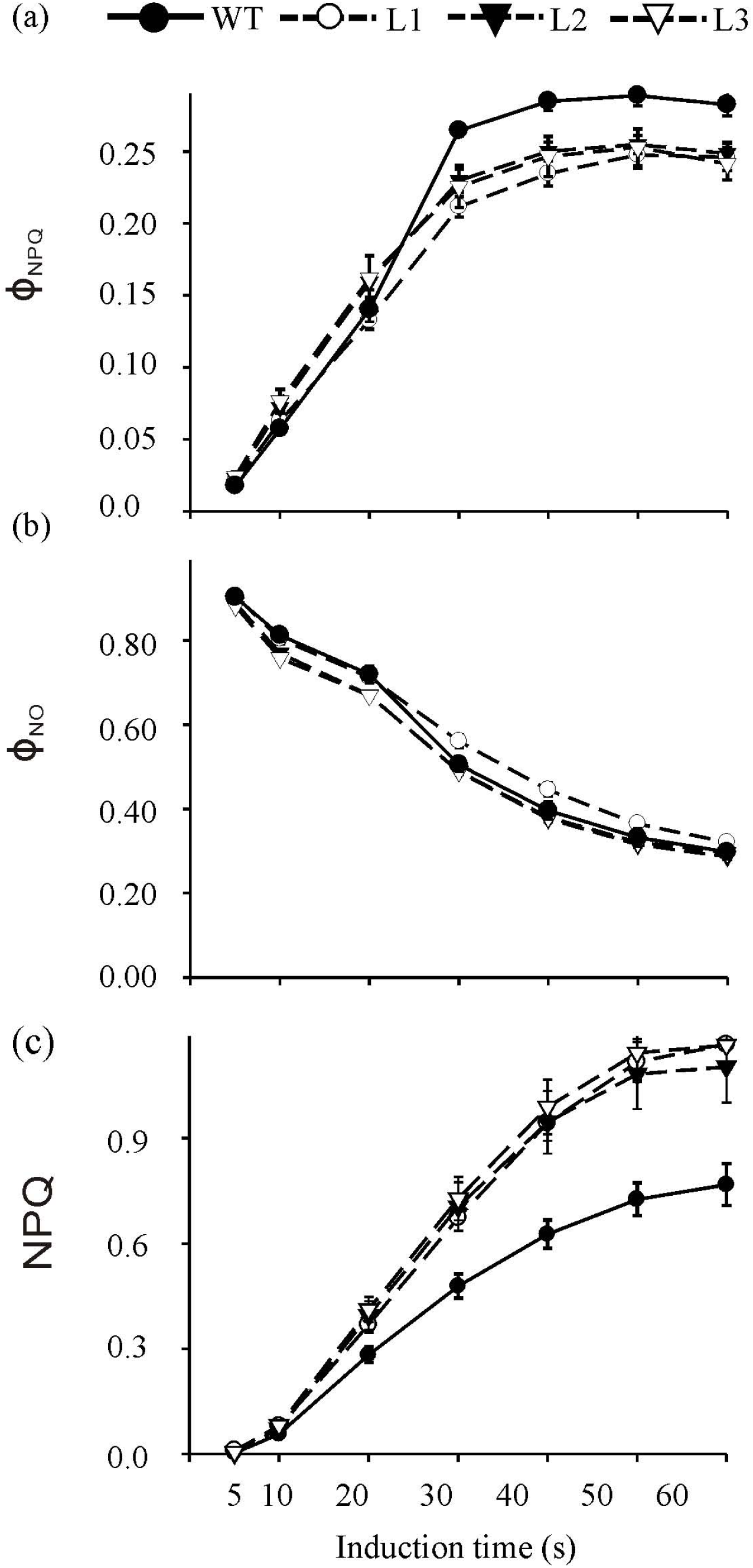
The effect of heterologously expressing *Flv* genes on non-photochemical quenching. The temporal behavior of dark-adapted leaves of six week old plants exposed to 160 μmol photons m^−2^ s^−1^ of actinic light with respect to (a) Φ_NPQ_ (the quantum yield of non-photochemical energy dissipation in PSII), (b) Φ_NO_ (the quantum yield of non-photochemical energy dissipation in PSII), (c) NPQ (non-photochemical quenching). L1-L3: lines harboring *Flv1/Flv3.* Data shown in the form *m*ean±SE (*n=*8).

### The effect of expressing Flv transgenes on leaf sugar, starch and amino acid content

The diurnal variation in the content of both soluble and insoluble sugars was determined in the leaves of WT and *Flv* transgenics (Figs 5 and S4). The leaves of plants harboring either *Flv1*/*Flv3* accumulated a significantly higher concentration of sucrose than did those of WT plants; in each line, the sucrose content increased gradually during the light period and fell during the dark period (Figure 5a). Leaf starch contents at the beginning of the light period did not vary between the genotypes, but the accumulation of starch during the day increased more rapidly (by as much as 1.7-fold) in the transgenics than in WT (Figure 5b). Glucose and fructose contents varied rather inconsistently (Figure S4). No clear genotypic differences were observed with respect to the accumulation of any of the amino acids following the plants’ exposure to 4 h of light (Table S1), but by the end of the light period (8 h of light), the increased pool of asparagine, aspartate, glutamine and alanine was observed in *Flv1*/*Flv3* transgenics with respect to WT (Table S2).

**Figure 5.**
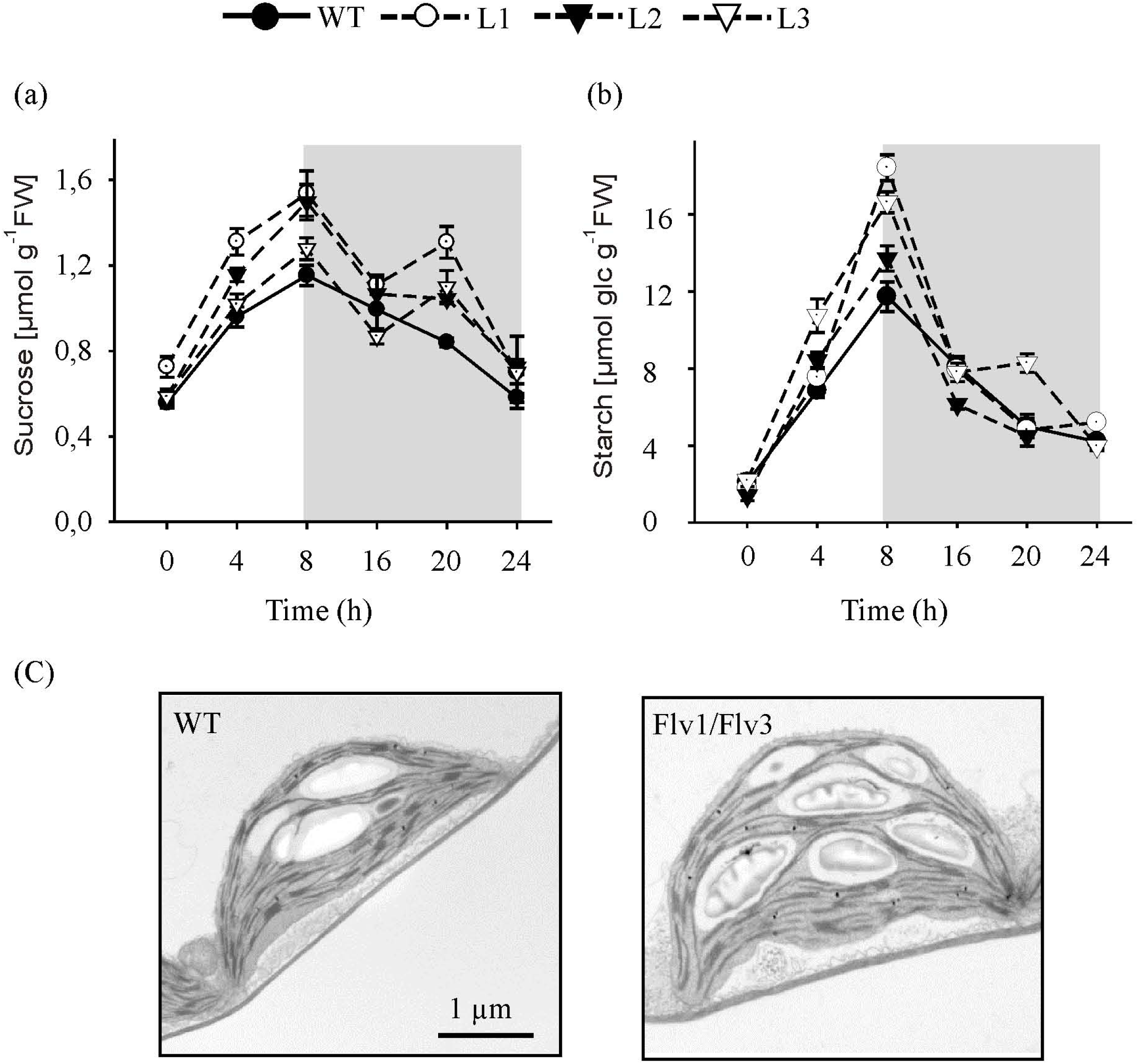
The effect of heterologously expressing *Flv* genes on diurnal variation in carbohydrate metabolism. Temporal variation in the rosette leaf of six week old plants exposed to 160 μmol photons m^−2^ s^−1^ of actinic light with respect to the content of (a) sucrose, (b) starch. (c) Representative images of starch granules present in leaves harvested after a 5 h exposure to light. The gray boxes indicate the dark period. Data shown in the form mean±SE (*n=*5).

### The effect of expressing Flv transgenes on starch granule size and number

Based on micrographs generated by transmission electron microscopy, it was apparent that while starch granule size was unaffected by the expression of *Flv*, their number was enhanced in the leaves of plants expressing *Flv1/Flv3* (Figure 5c).

### The effect of expressing Flv transgenes on the ATP levels in the leaf

The leaf content of ATP, ADP and AMP was measured both during the light period (after exposure to both 4 h and 8 h) and during the dark period (16 h). The ATP content of the *Flv1/Flv3* transgenics’ leaves was up to 1.5 fold higher than that of WT leaves, whether the leaves were sampled during the light or the dark period (Figure 6a). The ratio of ATP/ADP is higher at 4 h and 16 h in all the trangenics lines (Figure 6b). The leaf content of neither ADP nor AMP differed significantly between the *Flv1/Flv3* transgenics and WT (Table S3). The total adenylate content was not changed at all time points except L1 than that of WT plants (Figure 6c).

**Figure 6.**
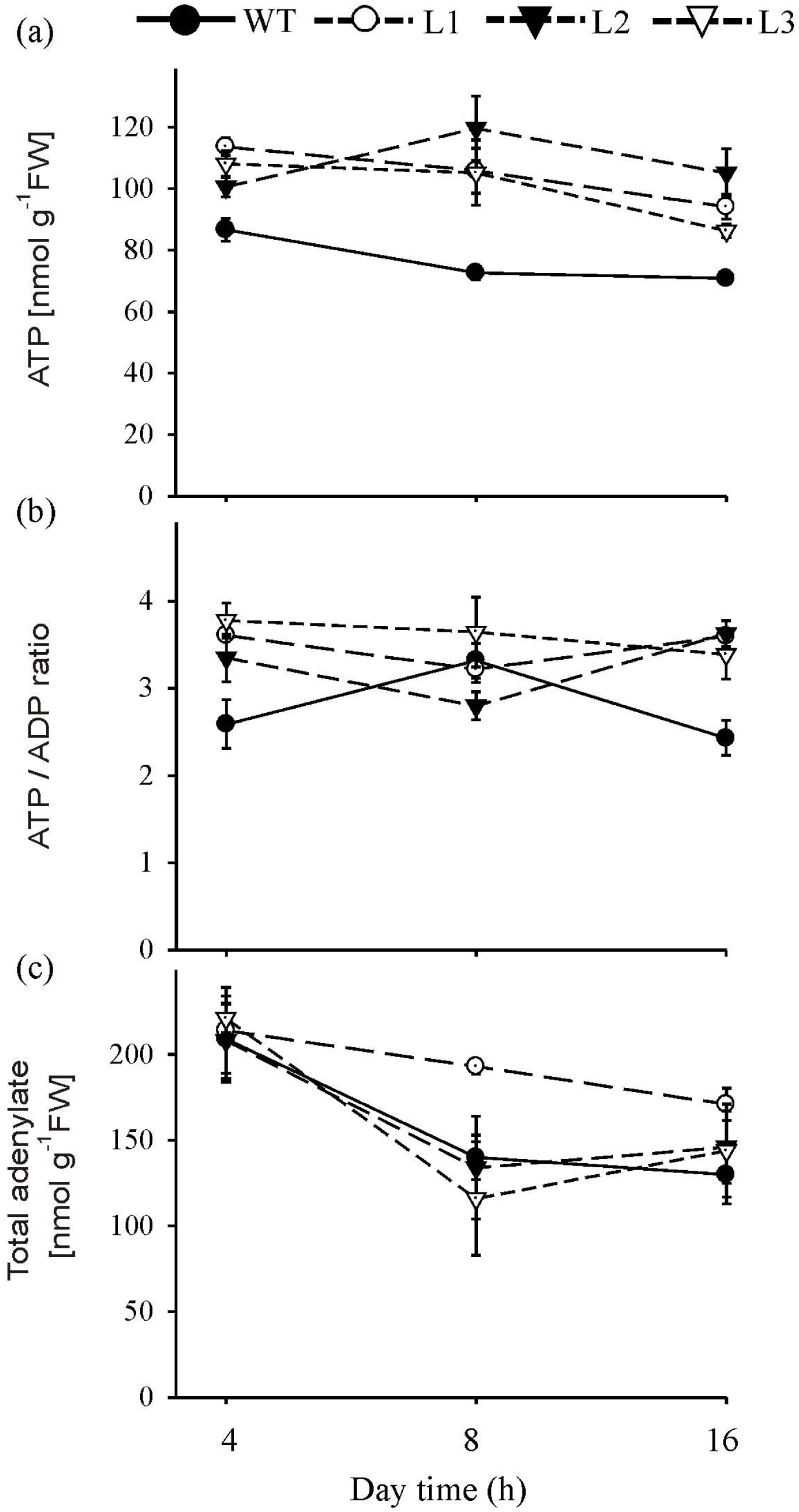
The effect of heterologously expressing *Flv* genes on the content of ATP, ATP/ADP ratio and total adenylate. Temporal variation in the rosette leaf of six week old plants exposed to 160 μmol photons m^−2^ s^−1^ of actinic light with respect to (a) the content of ATP, (b) ATP/ADP ratio and (c) total adenylate. L1-L3: lines harboring *Flv1/Flv3*. Data shown in the form mean±SE (*n=*5).

### The effect of expressing Flv transgenes on the leaf metabolome

The content of a number of metabolites was affected by the expression of *Flv* transgenes. After a 4 h exposure to light, the concentration of hexose phosphates was raised in all of the transgenic plants to a level significantly higher than that obtaining in WT leaves (Figure 7a); however, by the end of the light period, the hexose phosphate content was significantly below that in the WT leaf in all the transgenic plants (Figure 7b). When measured following the exposure of the plants to 4 h of light, the concentration of the starch precursor ADPGlc was elevated above the WT in all the transgenic plants (Figure 7c); the extent of the elevation was up to 1.4 fold for the *Flv1/Flv3* carriers (Figure 7c). When remeasured after an 8 h exposure to light, the level of ADPGlc remained statistically unchanged in the transgenics (the exceptions related to plants of lines L2) (Figure 7d). There was little evidence for any effect on the accumulation of citrate, or malate; significant elevations in the content of malate were recorded only in the leaves of line L2 assayed after a 4 h exposure to light, and in those of line L3 assayed after an 8 h exposure to light were statistically lower; at the 4 h time point, a modest increase was noted with respect to the concentration of citrate in the leaves of line L3 plants at 4 h and L1 plants at 8 h; (Figure S5a-d).

**Figure 7.**
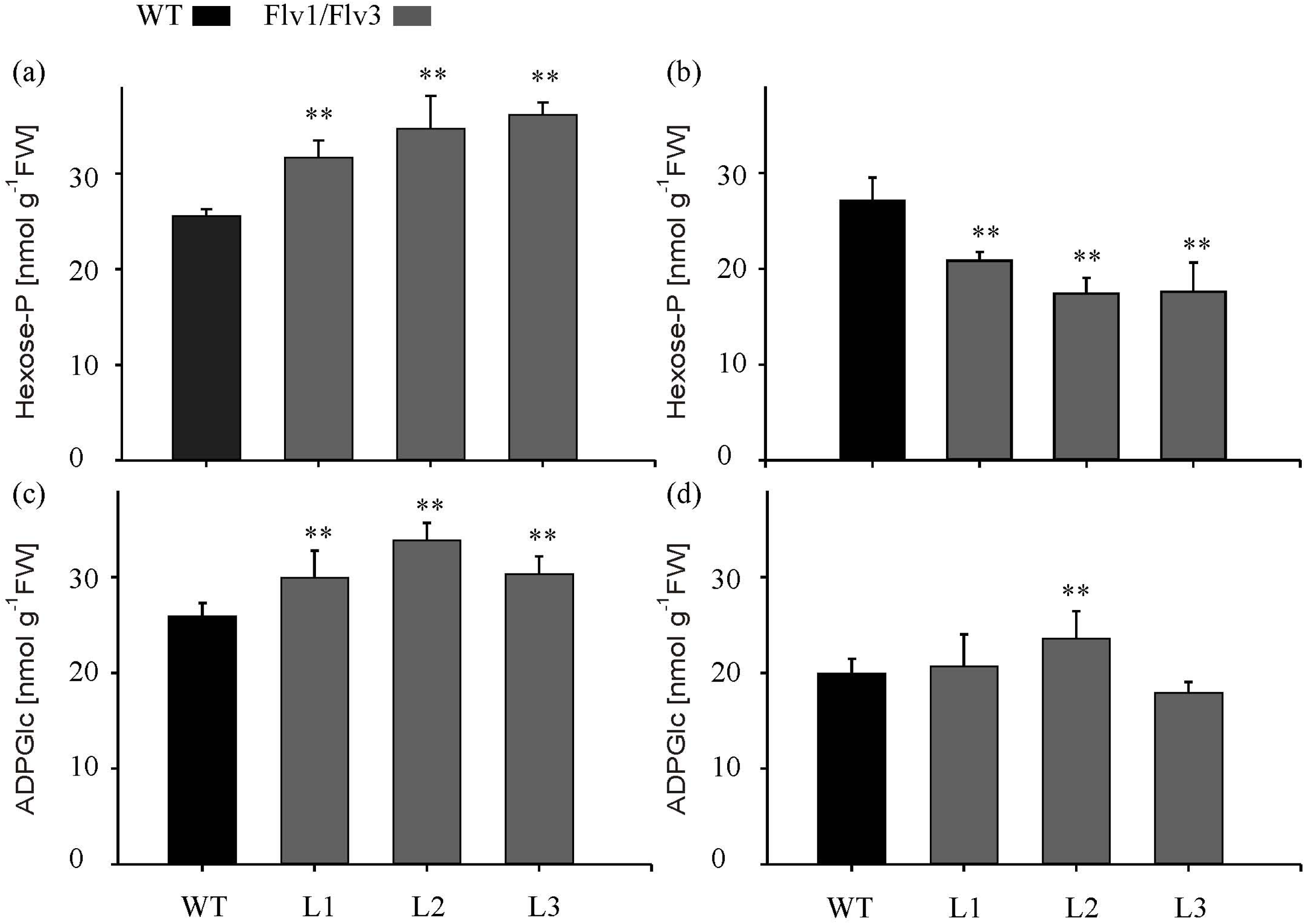
The effect of heterologously expressing *Flv* genes on the content of primary metabolites. The content in the rosette leaves of dark-adapted six week old plants exposed of (a,b) hexose phosphate, (c,d) ADPGlc, measured after exposure to (a,c) 4 h, (b,d) 8 h of light. L1-L3: lines harboring *Flv1/Flv3*. Data shown in the form mean±SE (*n=*5). **: means differ from the performance of WT at ≤0.01.

### The effect of expressing Flv transgenes on the content of antioxidants

The content of both the reduced and oxidized forms of glutathione were measured and their ratio was calculated. There was a significant shift towards the reduced form in L3 whereas oxidized form is statistically higher in line L2 of the transgenic leaves (Figure S6a,b), resulting in a reduction in the GSH/GSSG ratio of up to 2.5 fold (Figure S6c).

## Discussion

Photosynthesis is essential for the growth and development of plants, but the process is relatively inefficient since just 8–10% of the overall spectrum of solar radiation is used to convert CO_2_ to sugar, while only 2-4% of incident light energy is channeled into growth (Long et al., 2006; Zhu et al. 2010). The rest harvested solar energy is released as heat and fluorescence. Tapping the lost solar energy and avoiding uncontrolled ROS production potentially enhance photosynthesis and overall plant growth by establishing electron sinks in chloroplasts.

Here, the intention was to establish an additional electron sink downstream of PSI in *A. thaliana* by heterologously expressing either *Flv1* and/or *Flv3*. The observation was that plants harboring *Flv1/Flv3* were able to maintain PETC in a more oxidized state prior to the induction of the dark reaction of photosynthesis (Figure 3a). Furthermore, plants harboring *Flv1/Flv3* exhibited an improved performance with respect to ϕ_PSII_, qL (Figure 3b,c) and NPQ (Figure 4a,c); these outcomes imply that the transgenic plants were capable to enhancing the flow of electrons from PSII to PSI, as well as having an improved capacity to dissipate electron pressure in the PETC during the dark-to-light transition, and the ability to activate protective non-photochemical quenching. The implication is that Flvs can act as regulators of photosynthesis like CET and Mehler reaction, specifically avoiding the over-reduction of the PETC (Figure 3). Gómez et al. (2018) have observed that while the photosynthetic performance of tobacco plants heterologously expressing cyanobacterial *Flv1* and *Flv3* under steady-state light was similar to that of WT plants as observed in *Flv1/Flv3* Arabidopsis plants, their photosynthetic parameters - including electron transport and NPQ during the dark-to-light transition – implied an improved ability of their dark-adapted leaves to maintain their PETC in a more oxidized state and to enhance proton motive force. The present results have demonstrated that the targeting of Flv proteins to *A. thaliana* chloroplasts improved their ability to dissipate electrons at PSI as the Mehler reaction during early induction of photosynthesis.

In phototrophic organisms, AEF pathways are induced shortly after their exposure to light, thereby generating ATP to supply the Calvin-Benson cycle and to support photorespiration during the dark-to-light transition. As reported by Shikanai and Yamamoto (2017), under steady-state conditions, Flvs have poor access to either NADPH or Fd, due to activation of the Calvin-Benson cycle, but may regain functionality under highly reduced stromal conditions. Here, the growth of *A. thaliana* plants heterologously expressing *Flv* responded differentially to the intensity of light provided (Figure 1). Under low light intensity conditions, electron transport is typically limited by the availability of photons, so that there is no need of relief with respect to the electron pressure on the PETC. The latter becomes important as the light intensity increases, and the Flv system is able to dissipate surplus reducing equivalents as long as the light intensity does not become excessive (Figure 1). Indeed, plants look stressed at this irradiation levels, as revealed by the color of the leaves (presumably due to anthocyanin accumulation, a typical response to high light). Wild type plants exhibited light stress symptoms during early exposure but acclimatized later, a similar phenomenon observed in Flv-expressing plants. When exposed to a long day (16 h photoperiod), the presence of the *Flv* transgenes accelerated flowering (data not shown) and the plants biomass and seed yield potential were boosted (Figure 2b,c), illustrating the benefit of Flvs in conditions where surplus electrons are produced.

In chloroplasts, ATP is generated via the linear electron transport and cyclic electron transport pathways (DalCorso et al., 2008), while in the mitochondria; the respiratory electron transport chain makes an additional contribution for ATP during daylight hours (Bailleul et al., 2015). The level of ATP was higher in the leaves of the *Flv* transgenics than in those of WT after the plants’ exposure to either 4 h or 8 h of light, indicating that the Flvs were able to dissipate electrons at PSI, enhance linear electron flow and thereby establish the pH gradient required for ATP synthesis (Figure 6a). Here, we cannot exclude the possibility that heterologous expression of the *Flv*s provided an AEF able to function independently of the cyclic electron transport pathway and to contribute to the ATP demand, consequently enhancing growth. The adenylate pool is also an important regulator of plant metabolism (Geigenberger et al., 2010). In the *Flv* transgenics, an increased ATP level served to boost the level of activity in the Calvin-Benson cycle, which in turn helped to maintain a high level of hexose phosphate during the middle of the light period (Figure 7a,b). The hexose phosphates and ADPGlc (Figure 7a-d) accumulated by the *Flv* transgenics are most likely used to synthesize sucrose and starch during the day, serving to stimulate plant growth (Figure 5a,b). The absence of any effect of *Flv* expression on the level of the TCA cycle intermediates citrate and malate implies that these organic acids most likely play at best a minor role in determining biomass production (Figure S5a-d).

The manipulation of plastidial levels of adenylate kinase was shown in both potato - where it has succeeded in raising starch and amino acid content (Regierer et al., 2002) - and in *A. thaliana* - where it has been shown to boost the accumulation of amino acids and promoted plant growth (Carrari et al., 2005). In the *Flv* transgenics exposed to 4 h of light, however, there was no evidence for any significant alteration in the leaf’s amino acid content, suggesting that illuminated leaves converted most of their photoassimilate into starch (Table S1). In contrast, the content of each of aspartate, asparagine, glutamine and alanine was raised by the plant’s exposure to 8 h of light (Table S2), consistent with the observation that an increase in carbon availability enhances the assimilation of the nitrogen needed for protein synthesis and hence for the continuation of growth in the absence of light (Lawlor, 2002). Furthermore, the level of both sucrose and ATP were higher in the *Flv* transgenics than in WT, not only during the light period, but also during the dark period. According to Sharkey et al. (2004), the levels of both sucrose and ATP are highly dependent on carbon metabolism during the night, while both Sulpice et al. (2009) and Graf and Smith (2009) have shown that these levels constitute important determinants of biomass accumulation. It seems probable therefore that the growth advantage enjoyed by the *Flv* transgenics reflects their superior capacity to generate photoassimilate and ATP.

The ratio between the reduced and oxidized forms of glutathione was significantly lower in the leaf of all of the *Flv* transgenics than in WT leaf (Figure S6c). Glutathione makes a major contribution to balancing the redox status of the mitochondria and chloroplasts. Considering the pool of glutathione in all the organelles, the ratio GSH to GSSG was estimated to be 20:1 in control conditions in average (Mhamdi et al., 2010a). However, in the *Flv* transgenics, the level of GSH was lower than in WT, while that of GSSG was higher (Figure S6a,b). The presence of the Flvs likely inhibits the leakage of electrons from the PETC, thereby damping down the production of ROS. No convincing experimental evidence was obtained to show that level of ROS (H_2_O_2_) was lower in the leaves of the *Flv* transgenics than in WT leaves (data not shown). Overall, the effect of heterologously expressing *Flv* in *A. thaliana* appeared to impact the cellular redox homeostasis; the Flvs are unlikely to promote oxidative stress because surplus electrons are diverted to the alternative Flv sink (Figure 8).

**Figure 8.**
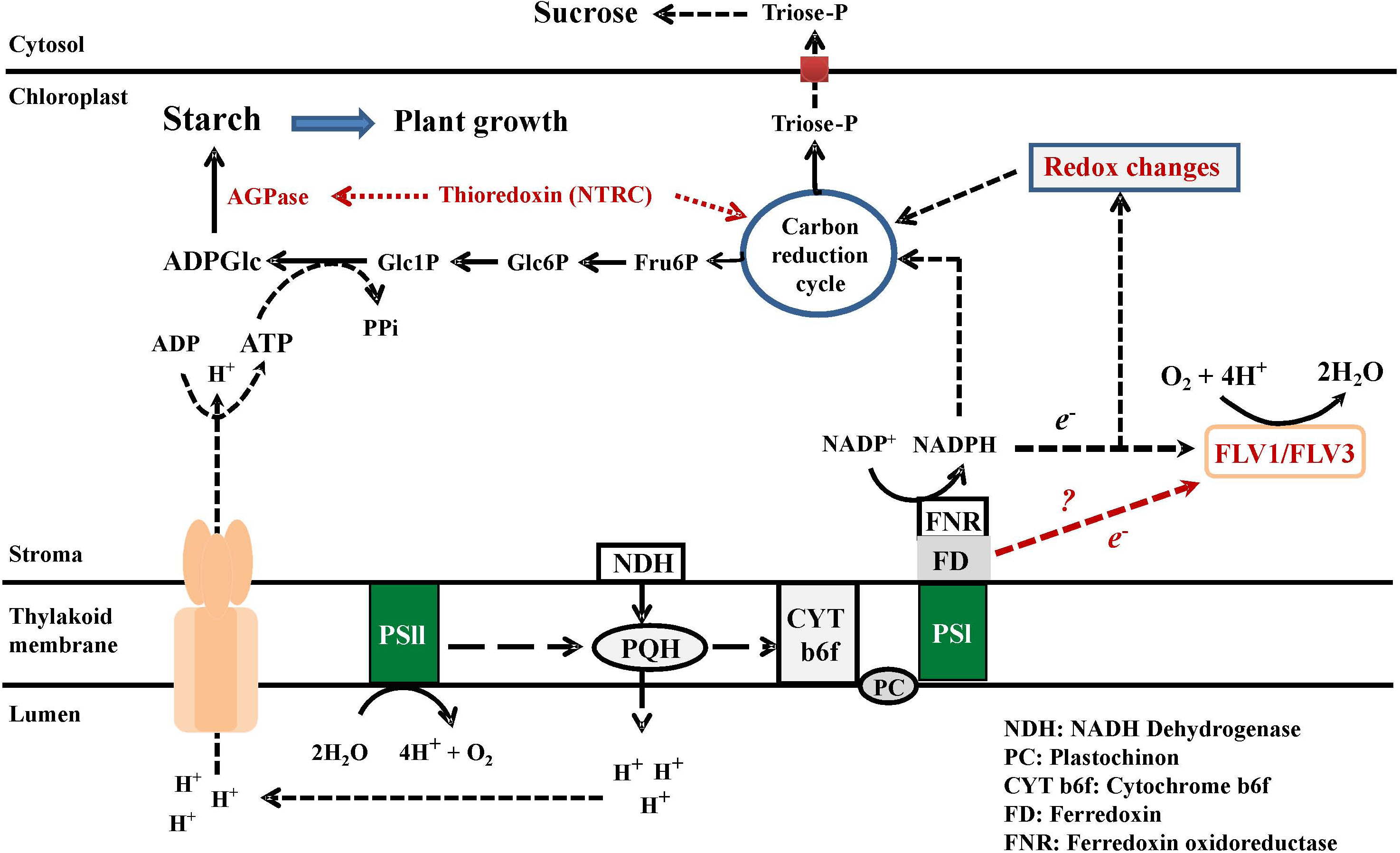
A model to explain the metabolic consequences of heterologously expressing *Flv* genes in the chloroplasts of *A. thaliana*. The presence of Flv proteins creates an electron sink and balances the surplus of the electron flow through PSI and PSII by delivering these electrons to oxygen, which is converted to water. The production of redox equivalents such as NADPH is maintained, resulting in the recycling of carbon through the Calvin-Benson cycle, which in turn generates an increased supply of the phosphorylated metabolites needed for starch synthesis. The energy required for this reaction is provided by ATP, which is synthesized in the PETC. The continuous flow of electrons results in the acidification of the lumen, which is the driving force for ATP synthesis. ATP is used for the conversion of Glc1P to ADPGlc via ADPGlc pyrophosphorylase. An increased availability of ADPGlc supports a higher level of starch synthesis; the starch accumulates when the leaf is exposed to light and is degraded during the dark phase.

## Conclusion

The present data have demonstrated that electron sinks are essential for the efficient functioning of the PETC and that they can be provided to an angiosperm such as *A. thaliana* by introducing genes encoding additional electron carriers such as cyanobacterial Flvs, either by expressing one *Flv* or by co-expressing two. The postulation is that the Flvs mediate the flow of electrons through PSI and PSII by delivering them to oxygen, which is converted to water without the production of ROS. In this way, the production of redox equivalents in the form of NADPH is maintained, allowing carbon to be recycled through the Calvin-Benson cycle. The latter generates the phosphorylated metabolites required for starch synthesis. The energy needed for this reaction is provided by ATP, which is produced via Flv-enabled electron flow and lumen acidification. ATP is also used for the conversion of Glc1P to ADPGlc, mediated by ADPGlc pyrophosphorylase activity. The promotion of ADPGlc finally results in an enhanced level of starch synthesis in leaves exposed to light, and the accumulated starch is metabolized during the dark phase, allowing for a continuous growth of the plant (Figure 8).

## Acknowledgments

We wish to thank Kirsten Hoffie, Marion Benecke, Melanie Ruff and Nicole Schäfer for excellent technical assistance in molecular and structural analysis at the IPK.

## Authors’ contribution

1. MRH and NC have made substantial contributions to conception and design, interpretation of the results and preparation of the manuscript

2. ST made all practical work and did the acquisition of data, or analysis and contributed to preparation of the manuscript

3. FS and MM have been involved in drafting and revising the manuscript, and preparation of the figures

4. TR, RG and AFL have been involved in preparation of microscopy figures (TR), calculation of photosynthetic parameters, interpretation of the data and revising the manuscript (RG, AFL)

5. NvW has contributed to the final revision and given the final approval of the manuscript

## Ethics approval and consent to participate

No animal research; not applicable.

No human subjects; not applicable.

## Supplementary Material

Figure S1. The vectors used to transform *A. thaliana.* (a) The design of the transgene constructs. The pea sequence encoding the transit peptide of ferredoxin oxidoreductase (TP, gray box) was fused to the N terminus of each *Flv* to drive expression in the chloroplast. (b) The abundance of *Flv* transcript in the transgenic plants. L1-L3: lines harboring *Flv1/Flv3*.

Figure S2. Subcellular localization of recombinant Flv. GFP activity in the chloroplasts of *N. benthamiana* transformed with in *GFP*-tagged *Flv1* and *Flv3*. The left panels show GFP fluorescence, the center ones chlorophyll autofluorescence and the right ones are merged images.

Figure S3. The effect of heterologously expressing *Flv* genes on the leaf content of NADPH. Measurements made after plants had been exposed to 8 h of light. L1-L3: lines harboring *Flv1/Flv3*. Data shown in the form mean±SE (*n=*5).

Figure S4. Diurnal variation in the sugar content of the rosette leaf in six week old plants heterologously expressing *Flv* genes. The content of (a) glucose, (b) fructose in transgenics harboring *Flv1/Flv3*. Data shown in the form mean±SE (*n=*5).

Figure S5. The effect of heterologously expressing *Flv* genes on the leaf content of organic acids. (a,c) Malate, (b,d) citrate. Measurements made after plants had been exposed to (a-c) 4 h, (d-f) 8 h of light. L1-L3: lines harboring *Flv1/Flv3*. Data shown in the form mean±SE (*n=*4 or 5). *,**: means differ from the performance of WT at, respectively, P≤0.05 and ≤0.01.

Figure S6. The effect of heterologously expressing *Flv* genes on the content of (a) GSH, (b) GSSG and (c) the ratio of reduced (GSH) to oxidized (GSSG) glutathione. Rosette leaves were sampled at the end of the light period (8 h photoperiod). Data shown in the form mean±SE (*n=*4 or 5).

Table S1. The contents of individual amino acids in WT and transgenic plants expressing *Flv1/Flv3*. Measurements taken from six week old plants which had been exposed to 4 h of light. Data shown in the form mean±SE (*n=*5).

Table S2. The content of amino acids in the leaf of WT and transgenic plants expressing *Flv1/Flv3*. Measurements taken from six week old plants which had been exposed to 8 h of light. Data shown in the form mean±SE (*n=*5). *,**: means differ from the performance of WT at, respectively, P≤0.05 and ≤0.01.

Table S3. The content of AMP and ADP in the leaf of WT and transgenic plants expressing *Flv1/Flv3.* Measurements taken from six week old plants which had been exposed to either 4 h, 8 h or 16 h of light. Data shown in the form mean±SE (*n=*5).

Table S4. List of gene specific primer sequences used for screening and quantitative RT-PCR analysis

---

**Table S1:** The concentrations of amino acids in WT and transgenic lines expressing Flv1/Flv3 genes. Measurements were carried out 4 h post illumination. Plants were six weeks old. Results are expressed as average and SE of 5 independent replicates. Significant differences are indicated by asterisks according to student’s t-test (\**P* ≤ 0.05 and \*\**P* ≤ 0.01).

| Amino acids [nmol x g<sup>-1</sup> FW] | WT | Flv1/Flv3 |  |  |
| --- | --- | --- | --- | --- |
|  |  | L1 | L2 | L3 |
| Asparagine | 369.8±21.3 | 317.8±26.3 * | 432.8±64.3* | 519.2±167.0* |
| Serine | 1197±109.2 | 1339±52.1 | 1685±286.7 | 1304±133.9 |
| Arginine | 22.3±1.0 | 20.5±2.1 * | 29.8±3.7 | 83.7±38.9* |
| Glycine | 55.8±45.0 | 13.9±1.8* | 16.2±5.5 | 24.4±19.2 |
| Glutamine | 903.1±94.7 | 842.7±47.9** | 1138±202.2** | 671.8±168.4** |
| Aspartate | 752.3±65.1 | 722.6±57.9** | 941.2±157.9** | 858.2±117.9** |
| Glutamate | 1742±131.6 | 1626±117.0 | 2035±310.9 | 1517±204.2 |
| Threonine | 649.9±65.7 | 628.7±42.6 | 878.8±131.9 | 640.4±94.3 |
| Alanine | 628.8±49.8 | 478.3±23.8** | 609.9±97.3** | 409.5±80.3** |
| GABA | 39.2±7.5 | 55.3±8.7 | 67.2±7.9 | 63.1±18.9 |
| Proline | 576.4±105.8 | 646.0±67.2 | 732.0±131.3 | 486.8±132.2 |
| Lysine | 19.2±1.7 | 23.2±2.7 | 27.0±4.2 | 28.7±1.1 |
| Valine | 63.4±4.7 | 58.9±6.1* | 67.8±9.6 | 91.0±17.3 |
| Isoleucine | 53.6±4.8 | 57.6±5.0 | 67.7±11.4 | 64.8±4.8 |
| Leucine | 17.7±1.6 | 18.6±1.9 | 23±4.0 | 17.4±5.4 |
| Phenylalanine | 14.6±1.4 | 16.9±2.4 | 19.1±3.8 | 17.8±5.0 |

**Table S2:** The concentrations of amino acids in WT and transgenic lines expressing Flv1/Flv3 genes. Measurements were carried out 8 h post illumination. Plants were six weeks old. Results are expressed as average and SE of 5 independent replicates.

| Amino acids [nmol x g<sup>-1</sup> FW] | WT | Flv1/Flv3 |  |  |
| --- | --- | --- | --- | --- |
|  |  | L1 | L2 | L3 |
| **Asparagine** | 265.1±6.7 | 325.5±23.1 | 332.5±19.5 | 405.8±58.9 |
| **Serine** | 991.3±17.4 | 948.4±76.9 | 966.7±13.4 | 945.3±22.3 |
| **Arginine** | 12.9±1.8 | 18.4±0.9 | 14.8±0.5 | 20.2±2.0 |
| **Glycine** | 53.2±4.8 | 86.6±12.8 | 76.0±19.6 | 96.3±25.3 |
| **Glutamine** | 1107±36.7 | 1502±77.4 | 1371±69.0 | 1439±33.4 |
| **Aspartate** | 570.9±15.6 | 712.8±51.3 | 615.9±19.3 | 650.1±34.9 |
| **Glutamate** | 1275±48.3 | 1194±57.6 | 1315±69.0 | 1300±40.7 |
| **Threonine** | 574.3±11.6 | 700.9±50.8 | 670.5±26.3 | 660.4±53.1 |
| **Alanine** | 437.2±25.4 | 684.9±32.8 | 602.4±35.6 | 551.1±17.7 |
| **GABA** | 23.9±2.7 | 25.3±2.1 | 32.1±4.7 | 32.7±4.9 |
| **Proline** | 402.3±24.5 | 588.1±59.9 | 656.7±64.9 | 551.2±26.0 |
| **Lysine** | 22.9±1.2 | 24.2±1.3 | 23.5±1.8 | 28.0±1.7 |
| **Valine** | 23.0±1.1 | 24.9±1.2 | 47.2±25.1 | 23.6±2.7 |
| **Isoleucine** | 55.1±1.0 | 68.4±4.9 | 61.0±2.7 | 60.2±5.2 |
| **Leucine** | 15.9±0.4 | 18.6±1.4 | 17.9±1.2 | 18.4±1.8 |
| **Phenylalanine** | 15.3±0.7 | 16.2±1.3 | 16.6±1.2 | 18.5±2.7 |

**Table S3:**
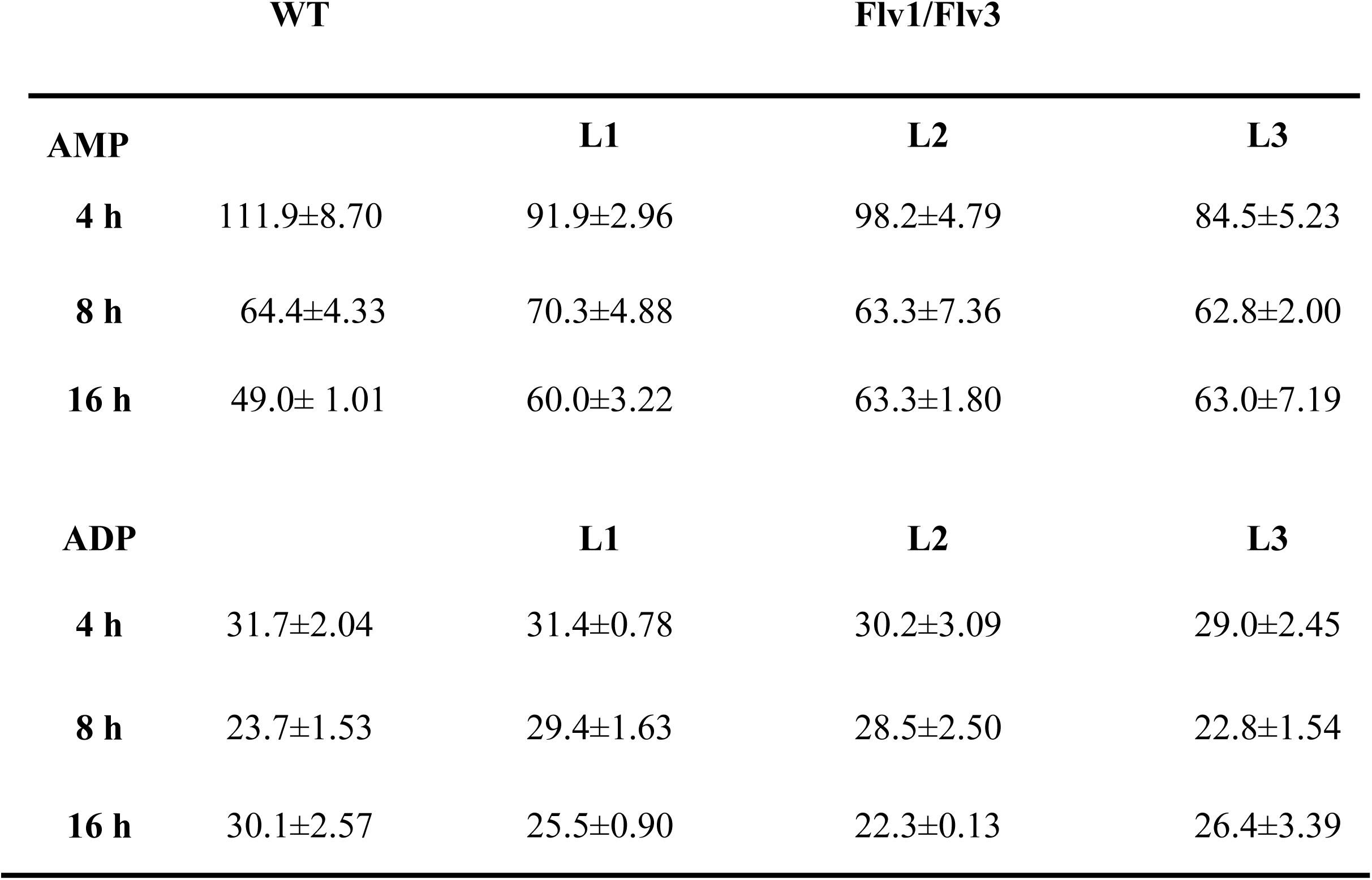
The Adenine nucleotide AMP, ADP concentrations in WT and transgenic lines expressing Flv1/Flv3 genes. Measurements were carried out at 4 h post illumination (2^nd^ time point), 8 h at the end of day light (3^rd^ time point) and 16 h dark period (4^th^ time point). Plants were six weeks old. Results are expressed as average and SE of 5 independent replicates.

**Table S4:**
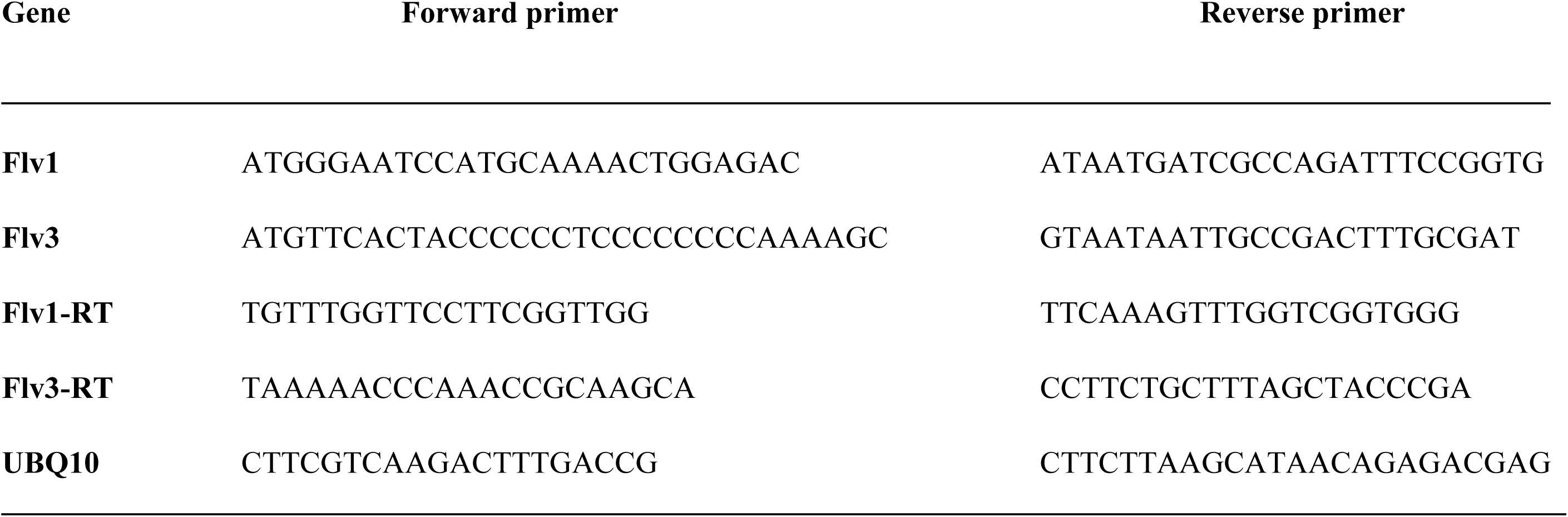
List of gene specific primer sequences used for screening and quantitative RT-PCR analysis

