## Supplementary figures and images for "Providing an additional electron sink by the introduction of cyanobacterial flavodiirons enhances the growth of *Arabidopsis thaliana* in varying light"

### Supplementary figures

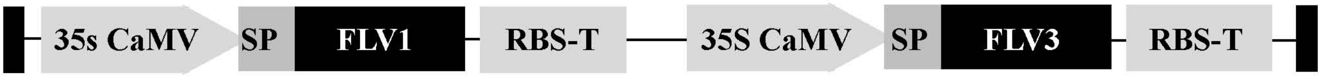

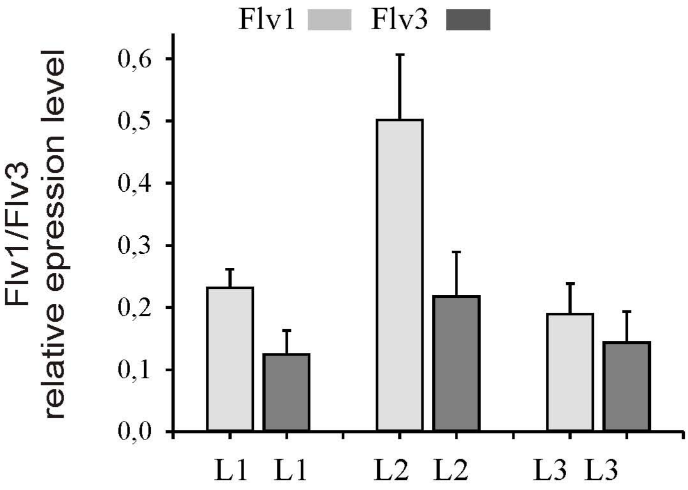

Figure S2

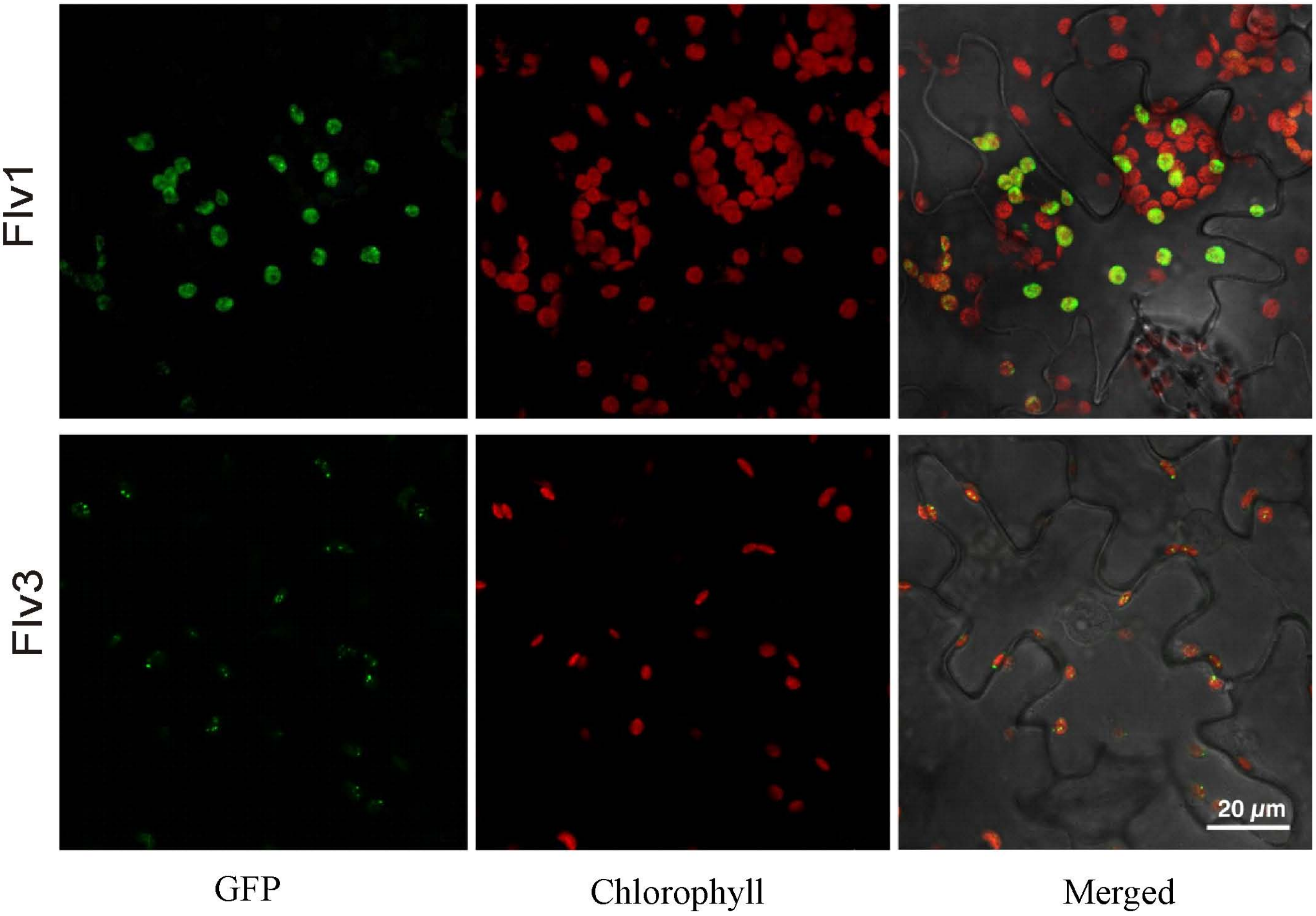

#### Figure S3

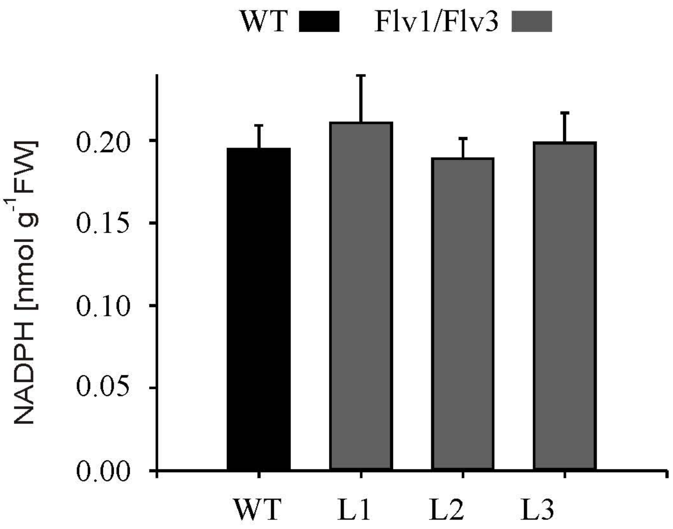

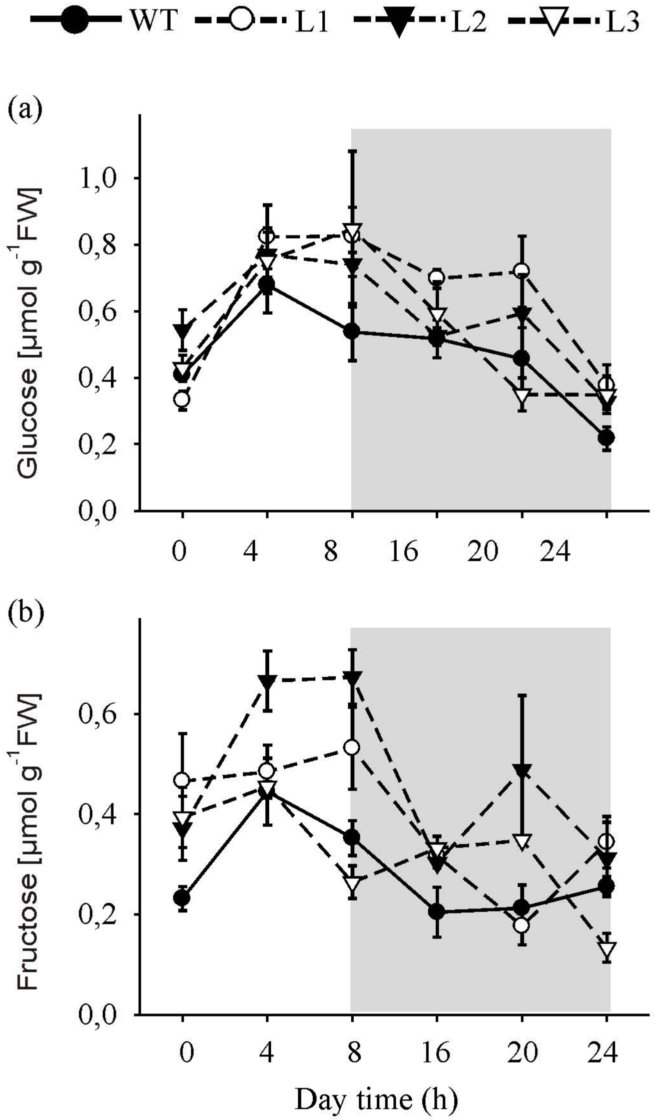

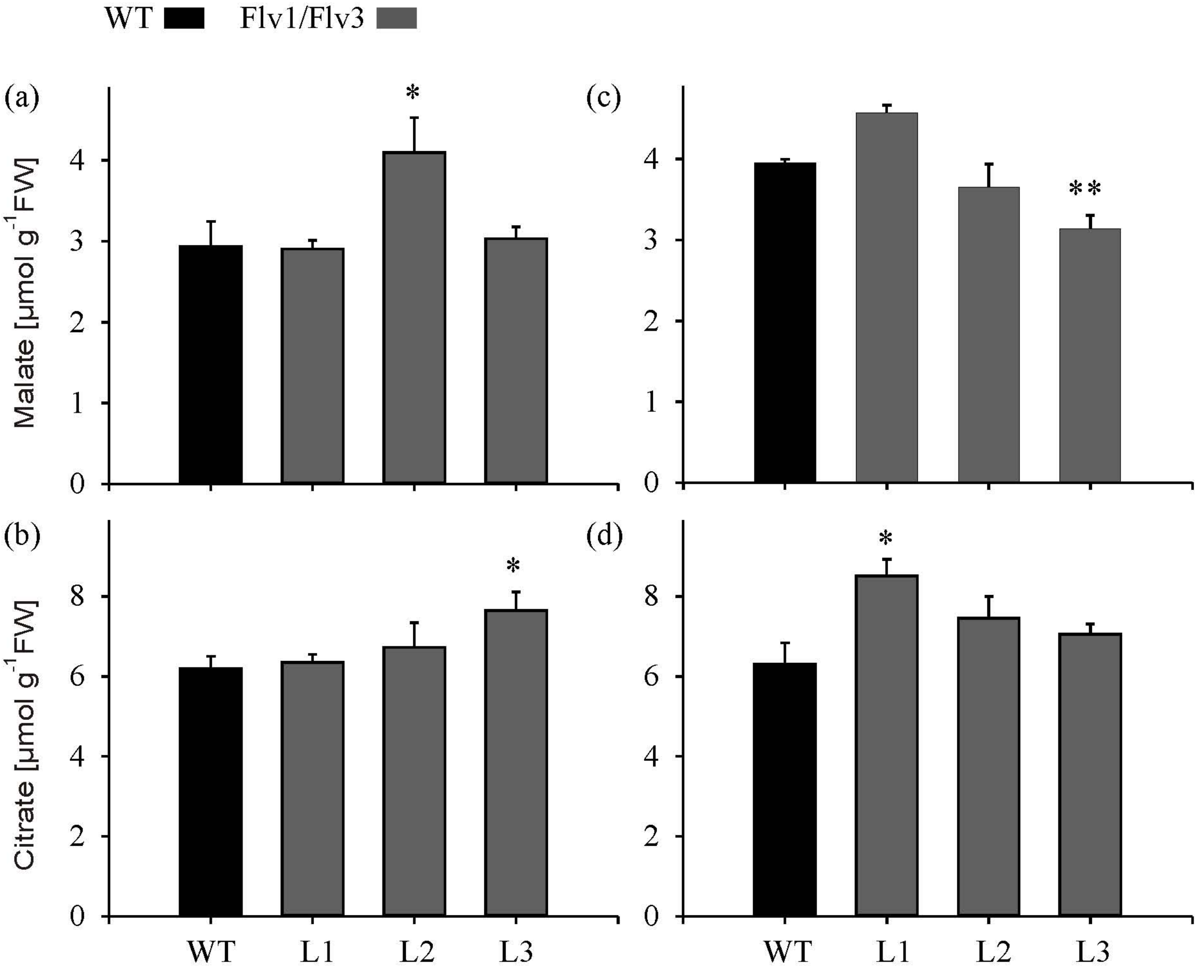

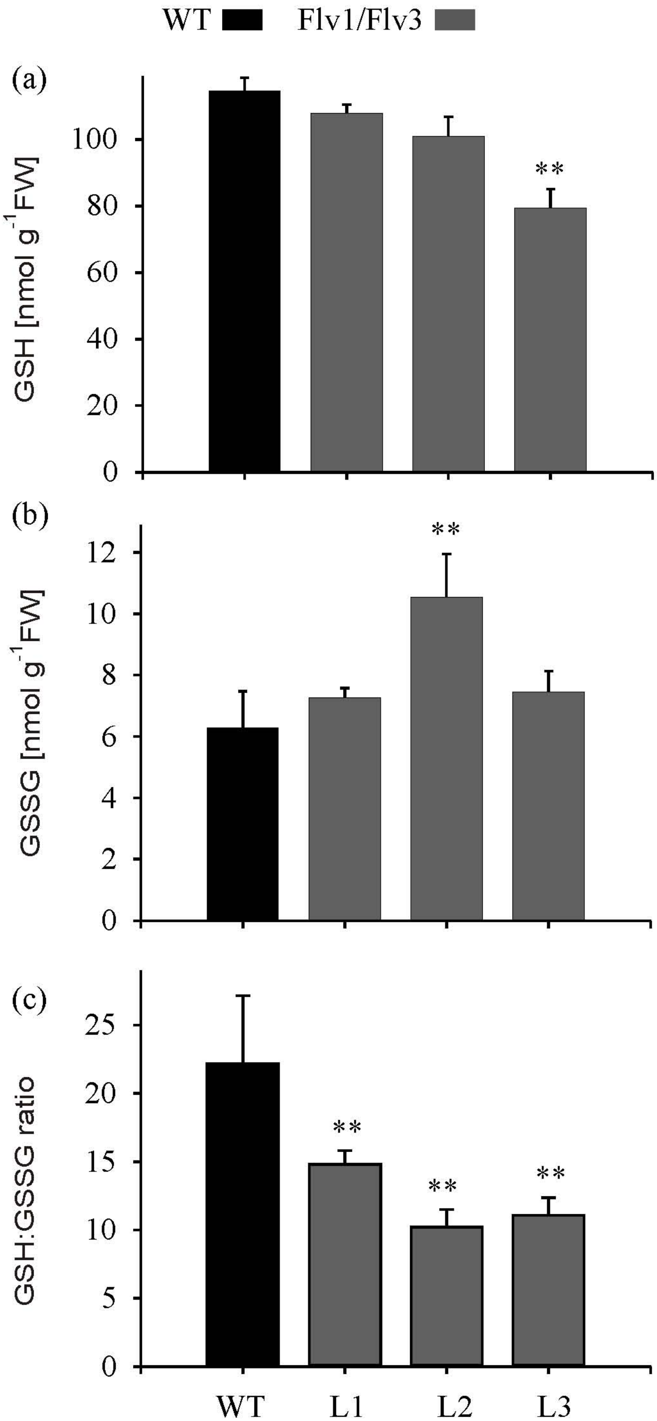
